## Supplementary Information for "Molecular Acclimation of *Halobacterium salinarum* to Halite Brine Inclusions"

#### 1 Supplementary Section 1: Halite Surface Contaminant Removal.

**Supplementary Table 1**: Summary of previously published protocols for the deactivation of microorganisms and removal of DNA from halite surfaces. It is important to note that none of these protocols were intended (or tested) for removal of proteins contaminating halite surfaces.

| Article | Dombrowski 1963<br>Radax 2001<br>Gruber 2003 | Norton & Grant 1988<br>Norton, McGenity, Grant 1993 | Stan-Lotter 1993<br>Denner 1994 | Rosenzweig 2000 | | Gramain 2011 | | | Sankaranarayanan 2011 | |
|---|---|---|---|---|---|---|---|---|---|---|
| **Samples** | natural halite | | | laboratory-grown halite: 96well plate naturally contaminated at the surface by an "unidentified white halophile" | | laboratory-grown halite (tested by coating halite surface with *Halococcus saccharoluticus* and/or *Halobacterium salinarum*) | | | natural halite from saline Valley California (immersion volume = 4 mL indicating relatively small crystal size) (tested by spiking halite surface with DNA; HV1 region of human mitochondrion) | |
| **Targets** | cell lysis | | | cell lysis (confirmed by CFU counts) | | cell lysis (confirmed by growth test) & DNA removal | | | DNA removal | |
| **Solutions** | NaCl unsatured | NaCl unsatured | NaCl unsatured | NaCl-saturated except for HCl | | NaCl-saturated except for HCl | | | NaCl-saturated except for HCl | |
| **Treatment & steps** | Bunsen burner | EtOH 100% -- 24h | EtOH 100% -- 6h | * 2 % SDS -- 1 min<br>* 0.02 % TA -- 1 min<br>________________<br>* 2 % SDS -- 1 min<br>* 0.02 % TA -- 1 min<br>*10 M NaOH -- 1 min<br>________________<br>* 2 % SDS -- 1 min<br>* 0.02 % TA -- 1 min<br>* 10 M NaOH -- 1min<br>* 10 N HCl -- 1 min | * 10 M NaOH -- 5 min<br>* saturated NaCl -- 2 min<br>* 10 N HCl -- 5 min<br>* brine saturated Na2CO3 -- 2 min | **A-**<br>* 10 M NaOH -- 5 min<br>* saturated NaCl -- 2 min<br>* 10 N HCl -- 5 min<br>* brine saturated Na2CO3 -- 2 min<br><br>**B-**<br>* 2 % SDS -- 1 min<br>* saturated NaCl -- 1 min<br>* 0.02 % TA -- 1 min<br>* saturated NaCl -- 1 min<br>* 10 M NaOH -- 1min<br>* saturated NaCl -- 1 min<br>* 10 N HCl -- 1 min | **C-**<br>* 70 % EtOH -- 1 min<br>* saturated NaCl -- 1 min<br>* 6 % NaOCl -- 1 min<br>* saturated NaCl -- 1 min<br><br>**D-**<br>* 70 % EtOH -- 3 min<br>* saturated NaCl -- 2 min<br>* 6 % NaOCl -- 3 min<br>* saturated NaCl -- 2 min<br>* 10 M NaOH -- 3 min<br>* saturated NaCl -- 2 min<br>* 10 N HCl -- 3 min<br>* saturated $Na_2CO_3$/NaCl -- 2 min | **E-**<br>* 6 % NaOCl -- 20 min<br>* saturated NaCl -- 5 min<br>* 10 M NaOH -- 20 min<br>* saturated NaCl -- 5 min<br>* 10 N HCl -- 20 min<br>* saturated Na2CO3/NaCl -- 20 min<br>* saturated NaCl -- 5 min<br>* 70 % EtOH -- 20 min<br>* saturated NaCl -- 5 min | # Al<br># Al-Bl<br># Al-Ac<br># Ac<br># Bl<br># Et<br># Bl-Et<br># Ac-Bl-Et<br># Ac-Et | # Al-Ac-Bl<br>* 10 M NaOH -- 15 min<br>* saturated NaCl -- 15 min<br>* 10 N HCl -- 15 min<br>* $Na_2CO_3$ -- 15 min<br>*saturated NaCl -- 15 min<br>* 6 % NaOCl -- 15 min<br>* saturated NaCl -- 15 min (x4)<br><br># Ac-Bl<br>* 10 N HCl -- 15 min<br>* $Na_2CO_3$ -- 15 min<br>* saturated NaCl -- 15 min<br>* 6 % NaOCl -- 15 min<br>* saturated NaCl -- 15 min (x4) |
| **Conclusions** | used | used | used | rejected | validated | rejected | rejected | validated | rejected | validated |

#### 2 Supplementary Section 2: Halophile Protein Extraction Protocol Summary for Bench Top Use

For all steps, sterilize solutions and materials by autoclave. All reagents should be analytical and HPLC grade. The use of glass materials including autoclaved glass Pasteur pipets is strongly recommended to avoid potential organic contamination from plastic polymers.

**Cell Collection and Lysis**

1. Sample preparation:
   A. **For liquid extraction:** Add sufficient volume of culture for 2 x $10^{10}$ cells to a glass centrifuge tube. Centrifuge 7 500 x g / 10 min / room temperature and discard supernatant.
   B. **For extraction from salt crystals:** Directly after removal of salt crystal surface bound contaminants using the NaOCl-NaOH-HCl active-spray surface organics method, add crystals to a glass centrifuge tube.
2. Add 5 mL of TRIzol reagent™ in each tube, ensure that crystals were fully immersed.
3. Crush crystal or cell pellet using autoclaved glass rod for approximately 5 min. Verify the absence of residual crystal fragments.
4. Vortex gently 30 s; incubate at 65 °C for 20 min.
5. Cooling tubes 5 min at room temperature (RT).

**RNA Fraction Removal**

6. Add 1 mL of 100 % chloroform in tube and parafilm.
7. Vortex gently 30 s; incubate at 5 min, RT
8. Centrifuge 10 000 x g, 20 min, 4 °C and discard very carefully upper colorless phase and dense middle phase.

**DNA Precipitation**

9. Add 1.5 mL of 100 % ethanol in tube and parafilm.
10. Vortex 3 s and incubate 3 min, RT.
11. Centrifuge 2 000 x g, 10 min, 4 °C
12. Collect carefully supernatant in a new 30 mL centrifuge tube.

**Protein Precipitation**

13. Add 7.5 mL 100 % isopropanol in tube and parafilm. Do not mix by vortex to avoid proteins adhering to the centrifuge tube wall.
14. Incubate 15 min, RT.
15. Centrifuge 10 000 x g, 20 min, 4 °C and very carefully discard supernatant.

doi: https://doi.org/10.1101/2022.09.16.508090

**Protein Pellet Desalting and Phenol Removal**

16. Add 5 mL of 0.3 M Guanidine-HCl in 95 % ethanol. Do not mix by vortex to avoid proteins adhering to the centrifuge tube wall.
17. Incubate 20 min, RT. If necessary, samples can be temporarily stored at -20°C before proceeding to the next step
18. Centrifuge 10 000 x g, 20 min, 4 °C and carefully discard supernatant.
19. Repeat steps 16-18.
20. Add 5 mL of 100 % ethanol, parafilm.
21. Incubate 20 min, RT.
22. Centrifuge 10 000 x g, 20 min, 4 °C and carefully discard supernatant.
23. Add 2 mL 100 % glacial acetone (-20 °C), parafilm.
24. Incubate 1h at -20 °C
25. Centrifuge 10 000 x g, 20 min, 4 °C and carefully discard supernatant.
26. Repeat steps 23-25.
27. Air drying protein pellet under hood (opened tubes in ice) until complete acetone evaporation.

**Protein Resuspension**

28. Gradually resuspend protein pellet with 2 mL 1 M $NaHCO_3$, 0.1 % sodium dodecylsulfate (SDS) at RT over 2 days. If the solution remains turbid, add more resuspension buffer.

**Protein Quantification Assay**

29. Quantitate resuspended proteins using the Pierce® BCA kit according manufacturer's instructions. After assay, aliquot proteins and stock at -20 °C for further analysis.

**Proteins Digestion**

30. Using a 100 µg aliquot of proteins from the previous step, reduced di-sulfur bonds using 2 mM of tris-(2-carboxyethyl) phosphine (TCEP) for 1h at 37 °C.
31. Alkylate cysteines with 5 mM iodoacetamide for 30 min at RT in the dark.
32. Digest proteins with 5 µg of Trypsin Gold (Promega) for 15 h at 37 °C with continuous agitation

**Peptide Desalting by Solid Phase Extraction (SPE)**

33. Use Sep-Pak C18 Plus cartridge Short 400 mg Sorbent (Waters) and elute peptides with 2 mL 70 % acetonitrile, 30 % ($H_2O$, 0.1 formic acid).
34. SpeedVac eluted peptides and resuspend them in 50 µL of 0.5 M TEAB

**Isobaric Labelling of Peptides**

35. Using 50 µg of initial proteins, labelled peptides with 8-plex iTRAQ® according manufacturer's instructions. Then desalting labelled peptides with SPE by repeating steps 33-34.

Resuspend labelled peptides in 2 % acetonitrile, 98 % (H2O, 0.1 % formic acid) using the appropriate volume for the desired peptide concentration. Stock at -20 °C until mass spectrometry analysis.

doi: https://doi.org/10.1101/2022.09.16.508090

#### 3 Supplementary Section 3: Protein Quantitation Using Modified BCA Assay.

Standard BCA protein assay allowed quantitation of protein concentrations between 125 and 2 000 µg with a microplate procedure using 1:20 sample:working reagent ratio. This protocol does not allow for detection of allowing low protein concentrations such as those obtained after halite surface cleaning. We therefore used modified concentrations of bovine serum albumin (BSA) standards for a working range between 25 and 200 µg proteins (see Supplementary Table 3-1), to establish the protein standard curve presented in supplementary Figure 3-1.

**Supplementary Table 3-1**: BSA standards concentrations for standard BCA and modified BCA assay.

| Standard procedure BSA concentration (µg/mL) | Modified procedure BSA concentration (µg/mL) |
|---|---|
| 2 000 | 200 |
| 1 500 | 175 |
| 1 000 | 150 |
| 750 | 125 |
| 500 | 100 |
| 250 | 75 |
| 125 | 50 |
| 25 | 25 |
| 0 | 0 |

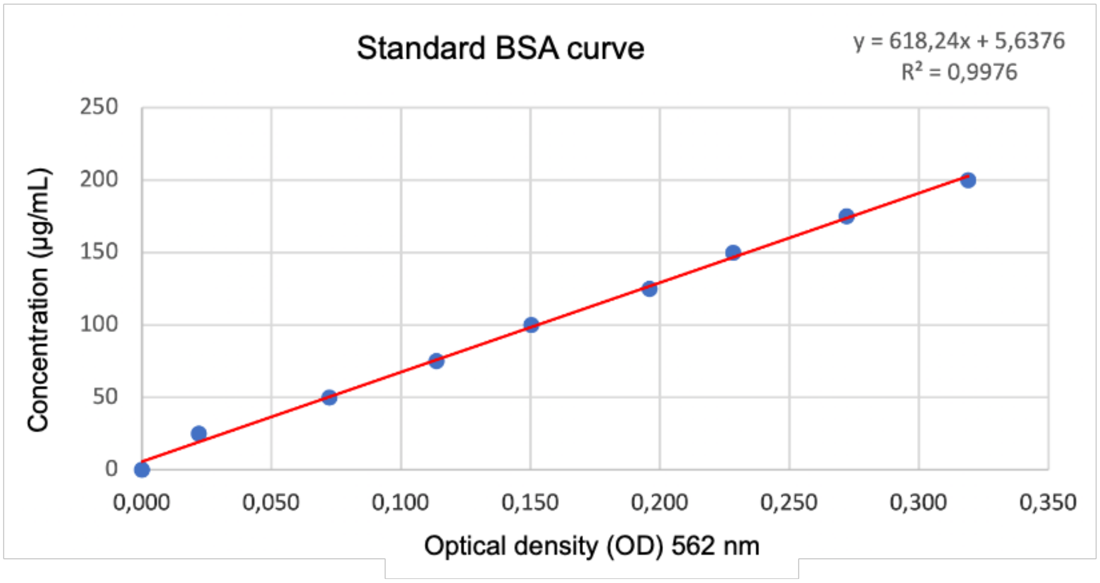


**Supplementary Figure 3-1**: Resulting modified BSA standard curve. Each concentration was done in triplicate and plotted as an average.

doi: https://doi.org/10.1101/2022.09.16.508090

### 4 Supplementary Section 4: Optimization of Protocol for Extraction and Analysis of Proteins from Halite Brine Inclusions.

This new TRIzol reagent™ based method for direct biomolecule extraction from halite crystals without prior dissolution step was optimized from the Kirkland *et al.,* 2006 procedure for liquid haloarchaeal cultures. As described in section 3.6 of the Materials and Methods, crystals were crushed directly in TRIzol Reagent™. The combined effects of the halite and the release of intracellular $K^+$ and $Cl^-$ from *H. salinarum* cells upon lysis resulted in high salt concentrations in the phenol phase. Kirkland et al., showed that salts were removed in the following isopropanol, guanidine HCl and acetone steps. To ensure that no salts remained in the final protein pellets (for example combined with traces of phenol) we used two washes with 95% ethanol, 0.3 M guanidine HCl. Then guanidine HCl traces were removed using 100% ethanol wash prior to additional washes in glacial acetone (-20 °C; x2).
However, the wash steps with glacial acetone rendered protein solubilization difficult. To resolve this problem, we tested four different solubilization solutions at room temperature and 4 °C. Buffers tested included $NaHCO_3$ 1M, 0.1 % SDS ("SDS Na"); $NH_4HCO_3$ 0.5 M, 1 % SDS ("SDS NH4"); $NH_4HCO_3$ 0.5 M, 8 M urea ("Urea NH4"); $NH_4HCO_3$ 0.5 M, urea 7 M, thiourea 2 M, CHAPS 4 % (w/v) ("Mix NH4"). Following solubilization and subsequent trypsin digest, LC-MS/MS (micro-injection) results (see Supplementary Figure 4-1) showed a higher number of peptides and proteins identified for urea, thiourea and CHAPS mix. However, the CHAPS buffer was ultimately eliminated due to its incapability with the BCA protein assay. Ammonium bicarbonate with 1 % SDS presented few proteins, which can be explained by partial digestion likely due to degradation of trypsin by high SDS content. Finally, while a comparison of ammonium bicarbonate with urea and sodium bicarbonate with SDS show similar results, the sodium bicarbonate buffer was ultimately chosen to avoid any interaction of ammonium ions during iTRAQ® labelling. Better results were obtained (slightly more peptides and proteins and less turbidity in sample tubes) at room temperature than at 4 ˚C.

Optimization of disulfide bonds reduction by TCEP was also performed to ensure optimal tryptic protein digestion. Two concentrations were tested as recommended by the supplier: 2 mM and 10 mM. The higher concentration resulted in fewer proteins than 2 mM concentration (data not shown), therefore 2 mM TCEP solution was used for reduction step.

Prior to LC-MS/MS injection, sample require cleaning and final desalting by solid phase extraction (SPE). We tested two commonly used cartridges: C18 and C8 (Sep-Pak Plus Short, 400 mg Sorbent). No differences were observed regarding number of peptides and proteins identified using either cartridge, and therefore C18 cartridges were selected for proteomics analyses.

The trypsin digests were analyzed by LC-MS/MS on an ultra-high-performance LC system (Ultimate 3000 RSLC, Thermo Scientific) connected to high-resolution electrospray ionization – quadrupole – time of flight (ESI-Q-TOF) mass spectrometer (Maxis II ETD, Bruker Daltonics). Separations were achieved on an Acclaim RSLC Polar Advantage II column (2.2 µm, 2.1 × 100 mm, Thermo Scientific) at a flow rate of 300 µL/min, using the following gradient of solvent A (ultra-pure water / 0.1 % formic acid) and solvent B (HPLC-MS grade acetonitrile / 0.08 % formic acid) over a total run time of 20 min linear increase from 2 % to 100 % B for 15 min, linear increase to 100 % B for 1 min, decrease to 2 % B for 1 min. The ESI-Q-TOF instrument was externally calibrated before each run using a sodium formate solution consisting of 10 mM sodium hydroxide in isopropanol / 0.2 % formic acid (1:1, v/v). Data-dependent LC-MS/MS data were acquired in positive ion mode in the mass range m/z 150 – 2200, using collision induced dissociation with collision energy calculated from m/z and charge states. The LC-MS/MS data were treated with Data Analysis 4.4 (Bruker Daltonics) and PEAKS software as previously explained in methods section.

doi: https://doi.org/10.1101/2022.09.16.508090

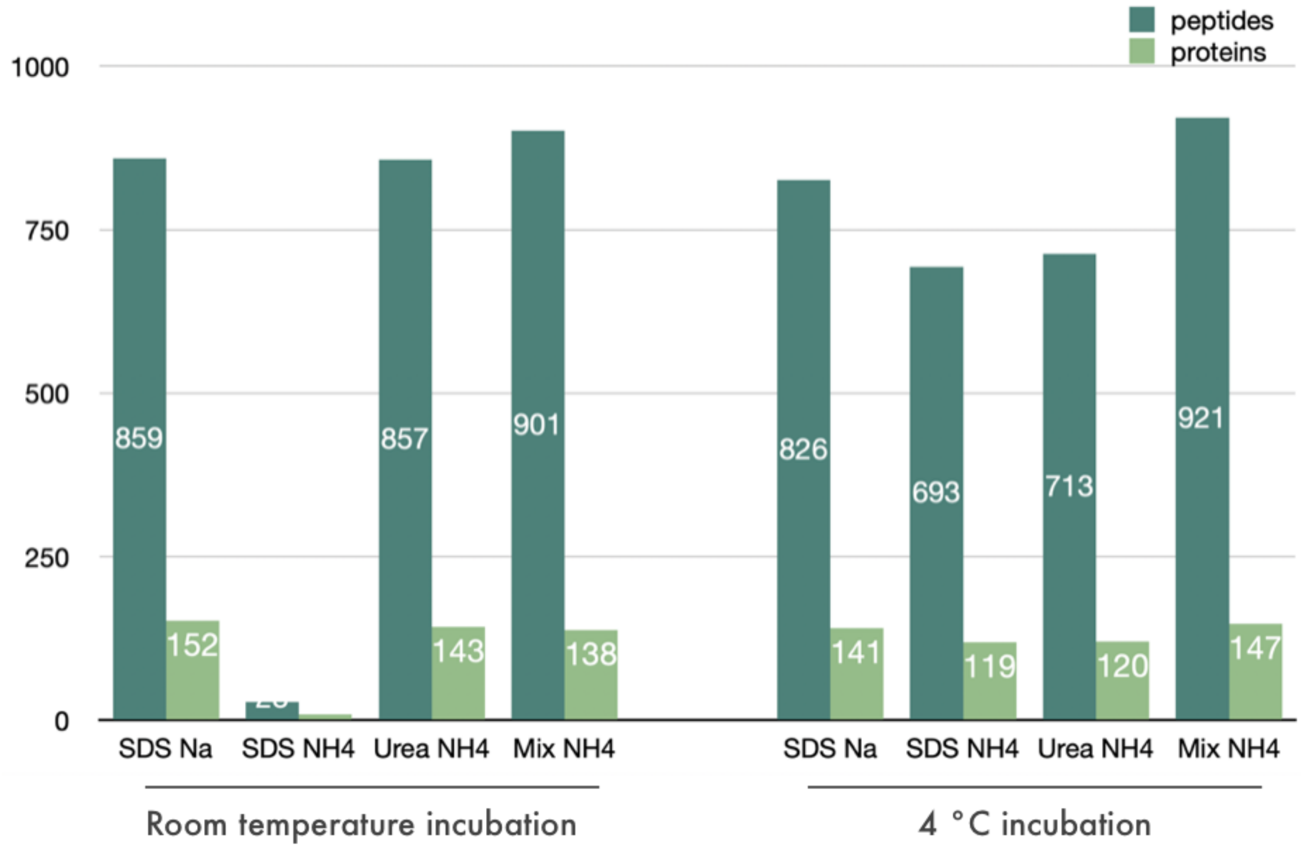


**Supplementary Figure 4-1**: Proteins and peptides identified by mass spectrometry for the four tested resuspension buffers: $NaHCO_3$ 1M, 0.1 % SDS (“SDS Na”); $NH_4HCO_3$ 0.5 M, 1 % SDS (“SDS $NH_4$”); $NH_4HCO_3$ 0.5 M, 8 M urea (“Urea $NH_4$”); $NH_4HCO_3$ 0.5 M, urea 7 M, thiourea 2 M, CHAPS 4 % (w/v) (“Mix $NH_4$”). Resuspension at room temperature and 4 °C were compared.

doi: https://doi.org/10.1101/2022.09.16.508090

#### 5 Supplementary Section 5: Comparison of Passive Baths and Active-spray Washes for Chemical Halite Surface Cleaning.

Chemical treatments for crystals surface cells and proteins removal were first tested as described in Sankaranarayanan *et al.*, 2011. Wash bath incubation times of 15 minutes completely dissolved the halite crystals, so baths were reduced to 5 min incubations. Proteins were then extracted from treated halite crystals using these passive bath washes. As seen in Suppl. Fig. 5-1, this protocol resulted in high levels of residual proteins. Instead, an active-spray procedure was tested for NaCl washes, resulting in near-total removal of proteins from halite surfaces when combined with NaOCl-NaOH-HCl treatments.

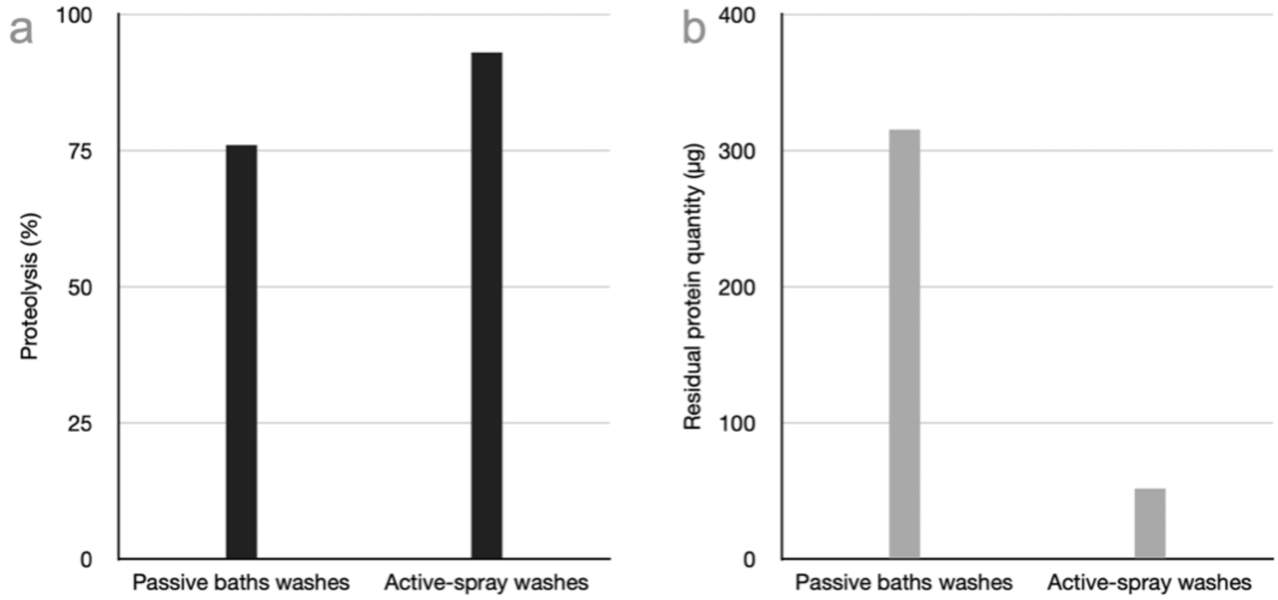


**Supplementary Figure 5-1:** Removal of surface-bound proteins using passive baths and active-spray washes during NaOCl-NaOH-HCl treatment on externally inoculated crystals.

#### 6 Supplementary Section 6: Complete Multi-Omics Results

Complete raw data sets for proteomics analyses are provided as separate .csv files (see Supplementary Tables 6-1, 6-2, 6-3 and 6-4).

Protein identification file **Supplementary Table 6-1** give PEAKS DB results without redundancies in proteins identifiers. Sheet 1 to 4 for liquids stationary samples (corresponding to Figure 4a), sheets 5 to 8 for brine inclusion extracts (corresponding to Figure 4b), sheets 9 and 10 for total core proteins of liquids stationary and brine inclusion samples respectively (Figure 4a,b). Sheets 11 and 12 summarized specific proteins for liquids stationary and brine inclusion samples. Sheet 13 corresponds to shared core proteins between both conditions with multiple proteins identifiers and correlated descriptions.

KEEG blastKOALA matched K numbers given in **Supplementary Table 6-2** were subdivided as sheet 1 for shared proteins between liquid and brine inclusions, sheet 2 for specific proteins of brine inclusion extracts, sheet 3 for specific proteins of liquid stationary extracts, sheet 4 corresponding to quantified proteins showing up-regulation and sheet 5 for quantified proteins showing down-regulation from brine inclusions.

KEGG pathway search BRITE results were given in **Supplementary Table 6-3**. Results for specific proteins identified in brine inclusion extracts (sheet 1), specific proteins identified in liquids stationary extracts (sheet 2) and those shared between both conditions (sheet 3). Then results for proteins showing significative up-regulation (sheet 4) and down- regulation (sheet 5) from brine inclusions.

**Supplementary Table 6-4** correspond to raw PEAKS Q quantitation results for iTRAQ sample injections for all eight samples (four replicates each liquid stationary phase extractions and halite brine inclusions extractions); first injection (replicate 1, sheet 1), second injection (replicate 2, sheet 2) and third injection (replicate 3, sheet 3) with redundancies in protein identifiers removed. Sheet 4 recapitulates the 68 proteins identified after pool of tree replicates results with multiple proteins identifiers and correlated descriptions.

**Supplementary Table 6-5** summarizes proteins identified and quantified for targeted pathways. These table include identified proteins shared by the four stationary and four brine inclusion extracts, proteins partially shared by the eight samples, and proteins with significant differential expression in brine extract compared to stationary cells.

RNA extractions were performed for both liquid cultures and halite (internally and externally inoculated) samples. A summary of RNA quality assays with BioAnalyzer is shown in **Supplementary Figure 6-2.**

doi: https://doi.org/10.1101/2022.09.16.508090 

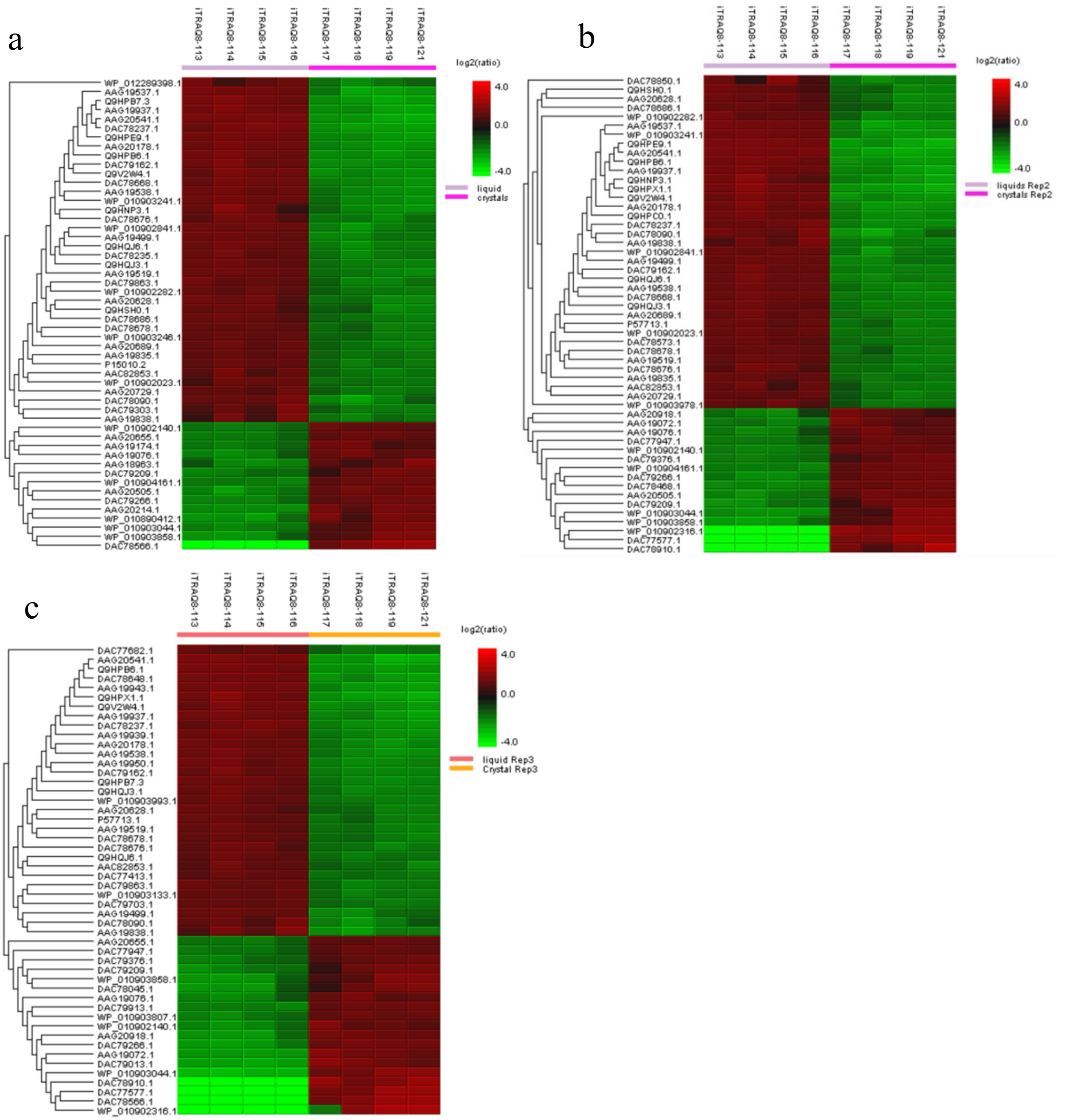


**Supplementary Figure 6-1**: Heatmaps for identified proteins showing differential expression in brine inclusions extracts ("crystals") compared to liquid stationary-phase cultures ("liquid") for the first (a), second (b) and third (c) injection replicates.

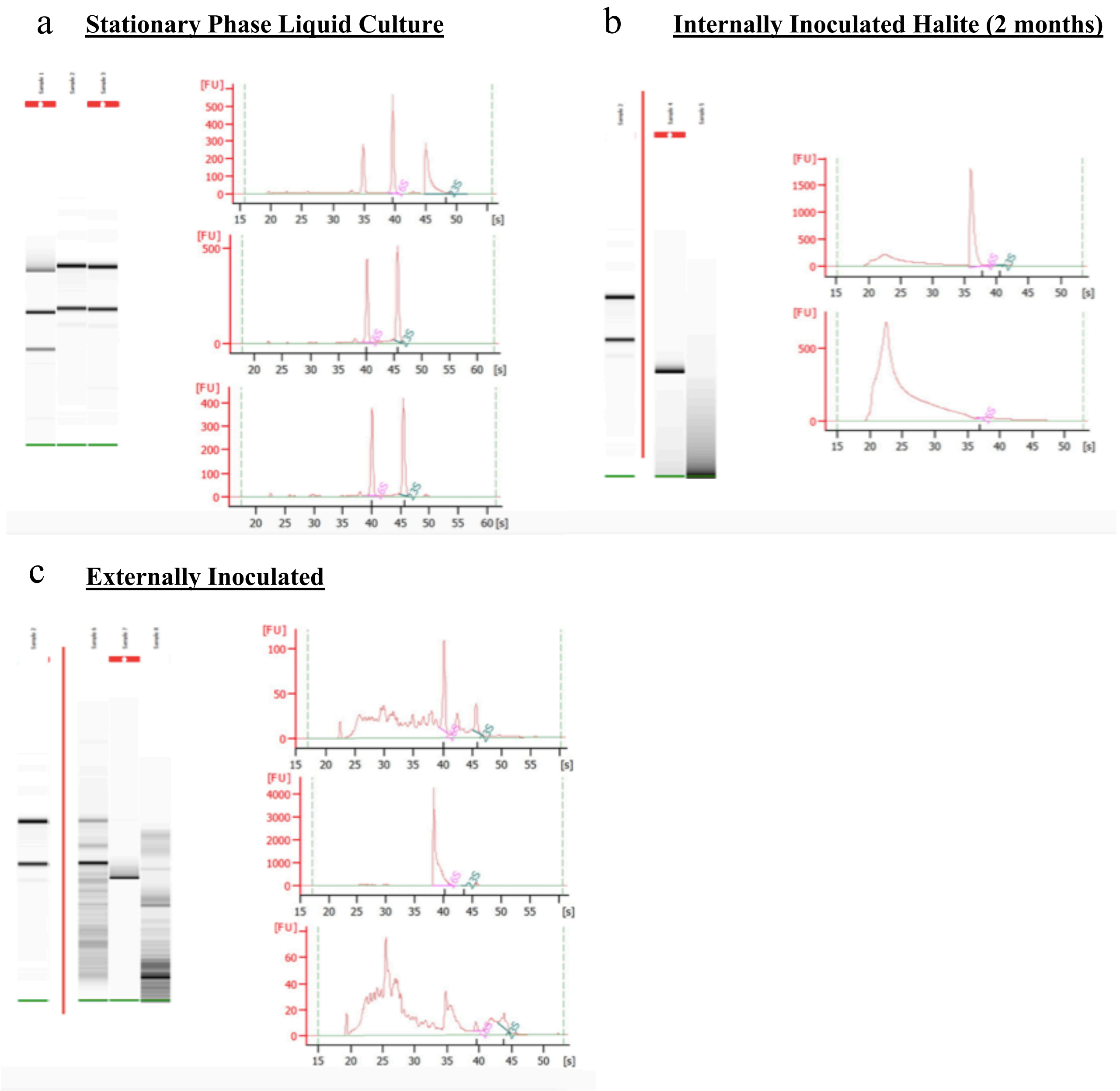


**Supplementary figure 6-2**: BioAnalyzer RNA quality results after TRIzol-based extraction and purification. RNA from (a) cells from liquid stationary phase cultures, (b) internally inoculated laboratory-grown halite after two months of incubation at 37 °C, and (c) externally inoculated laboratory-grown halite (no incubation period after drying).

#### 7 Supplementary Section 7: The Special Case of Transmembrane Peptides.

TRIzol Reagent™-base protein extraction seems unsuitable for transmembrane peptides. As shown below for ArcD arginine-ornithine antiporter example (Supplementary Figure 7-1), none of the transmembrane regions of the ArcD protein were identified in this study, whereas regions extending outside the membrane were detected by LC-MS/MS.

MVEFEPRSYEDFDPEKRPSFGQ**ALLPIAGMITFLAVGIVLLGL**DAQM**PLLWGIAFTGVIARYGWGYTW**DELFDGISNS**IVMGLGAIFILFIIY**
**MLIASW**VDAGTIPFIMYWGLEF**LTPAVFVPLAALLSFVVATAI**GSSWTTAGSLG**IALVGIGSGLGIPAPLTAGAI**LSGVYMGDKQSPLSDTTLL
ASGVSDVDLWDHVRG**MFPNTIIVGVISLALYAVLG**LMAETGGTGGTGAEVAQIQGGLA**GTYTLSVLVLLPLVITFGLAI**KG**YPALPSLGAGV**
**FSAAGVSILI**QGRGFAEAWQIIYSGTGPKTGVDLVNNLLSTGG**LEGSIWVITIVFGALSIGGIL**EATGVLSVIAHNTAKAVD**SVGGATLVTALGP**
**LVINALTA**DQYMSIVIPGMTFRDLNDEYDLDGTSLSRTLEETGTVSEPMIPWNSGGVFMASALGVP**VLSYLPYYFVGILSPILVVIM**GFTGWK
MYMKDPEESPEESADTAA

**Supplementary Figure 7-1**: Arginine-ornithine antiporter ArcD amino acid sequence with transmembrane peptides in blue and peptides identified in samples 1, 2 and 3 from liquid stationary-phase culture extracts in yellow.
